# Inference of multi-enhancer interactions in T lymphocytes using Hi-Cociety

**DOI:** 10.1101/2025.06.12.659372

**Authors:** Sora Yoon, Golnaz Vahedi

## Abstract

Three-dimensional (3D) enhancer communities are key regulators of gene expression, shaping cell fate decisions and contributing to disease pathogenesis. Assays such as H3K27ac HiChIP have been used to map enhancer–enhancer interactions and define enhancer communities; however, their reliance on antibody-based enrichment restricts scalability and cross-cell-type applicability. In contrast, Hi-C provides an unbiased, genome-wide view of chromatin architecture but lacks direct annotation of regulatory elements, limiting its utility for enhancer-focused analyses. To bridge this gap, we introduce Hi-Cociety—a graph-based computational framework and accompanying R package that infers 3D enhancer communities directly from Hi-C data, without relying on histone modification or chromatin accessibility measurements. Hi-Cociety constructs a network of significant interactions and applies clustering algorithms to define chromatin interaction modules. Applying Hi-Cociety to Hi-C measurements in T lymphocytes, we identified highly connected modules enriched for active transcription, chromatin accessibility, and histone acetylation. Notably, modules identified in T cells pinpoint critical genes central to T cell biology. Hi-Cociety also detects cell-type-specific differences in chromatin organization, highlighting dynamic regulatory rewiring across T cell states. Our findings underscore the importance of network properties— connectivity, transitivity, and centrality—in shaping gene regulation through 3D genome organization. Hi-Cociety provides a scalable and versatile tool for mapping enhancer communities at scale, advancing our understanding of immune cell identity and the regulatory logic encoded in 3D chromatin structure.

## Introduction

Three-dimensional (3D) enhancer communities are spatially clustered groups of enhancers that engage in coordinated regulation of gene expression^1–4^. These enhancer hubs play a critical role in orchestrating the expression of key genes involved in cell fate decisions by facilitating long-range chromatin interactions that enhance transcriptional output. By bringing multiple enhancers into close proximity with their target genes, 3D enhancer communities establish a regulatory architecture that integrates multiple signaling inputs, ensures robustness in gene expression, and enables precise control of lineage-specific transcriptional programs. Disruptions or hijacking of these enhancer communities can lead to aberrant gene expression, ultimately contributing to developmental disorders and diseases such as cancer^5^. Enhancer communities have been primarily defined using the H3K27ac HiChIP assay, which simultaneously maps enhancer-enhancer interactions. Graph-based methods are straightforward strategies to infer enhancer interactions using H3K27ac HiChIP^3–5^. Despite the importance of H3K27ac HiChIP in enhancer-enhancer mapping, it can be considered as a biased assay since it relies on interactions of antibody-pull down genomic regions. While there are around 1000 publicly available H3K27ac HiChIP data across different species and cell types, more than 20,000 publicly available Hi-C data can be accessed in public domains such as NCBI:GEO or 4DN data center. However, Hi-C lacks the direct enhancer annotation provided by histone modifications, necessitating computational approaches to infer 3D enhancer communities from these datasets. Developing a robust computational framework to extract enhancer hubs from Hi-C data would significantly expand our ability to study gene regulatory architecture at an unprecedented scale, providing deeper insights into the role of 3D chromatin organization in cell fate determination and disease mechanisms. We introduce Hi-Cociety, a scalable and versatile tool for identifying 3D enhancer communities from Hi-C and Micro-C datasets. By enabling the analysis of enhancer architecture across the wealth of publicly available Hi-C data, Hi-Cociety opens new opportunities to systematically investigate gene regulatory networks across diverse cell types and disease contexts. We also provide a comprehensive list of highly connected enhancer modules in T lymphocytes for the broader research community, which is freely available in Hi-Cociety website: https://sorayoon.shinyapps.io/HiCociety/.

## Results

### Hi-Cociety extracts chromatin interaction network modules from Hi-C data

We developed Hi-Cociety to infer modules of highly interacting loci leveraging Hi-C or other chromatin conformation capture assays. Hi-Cociety identifies significant chromatin interactions and represents them as a graph, enabling the reconstruction of interaction modules across each chromosome. Considering the noisy nature of inter-chromosomal interactions, only intra-chromosomal interaction are considered by Hi-Cociety. Hi-Cociety also reports differential modules exhibiting significant differences between two experimental conditions.

Hi-Cociety comprises multiple statistical components (Figure 1a). At its core, it models observed contact frequencies using a negative binomial distribution. For each linear genomic distance—ranging from the resolution of Hi-C contact frequency matrix (e.g., 10 kb) to 2 Mbp in fixed intervals—we estimate the distribution parameters: **μ** (mean) and **α** (dispersion). These parameters are used to define the expected contact frequency distribution for each 1D distance and each chromosome separately. We then compute a *P*-value for each pair of genomic loci with an observed contact frequency and identify statistically significant chromatin interactions. An additional filtering step involves removing chromatin interactions located in ‘contact-desert’ regions. For each significant chromatin contact (i.e., a pair of genomic loci), a padding of 5 pixels is applied around the genomic pair. If the average contact frequency within this extended region falls below a predefined threshold (N), the interaction is excluded from further analysis. The genomic pairs that pass all filtering steps serve as the network’s nodes and interactions with significant *P*-values serve as the network’s edges. A representative example of significant interactions resulted from the combination of these two filtering steps is shown at the *Rab51b* locus in large pre-B cells^6^ (Figure S1). Thus, Hi-Cociety infers significant chromatin interaction from Hi-C without relying on existing loop-calling methods.

**Figure 1.**
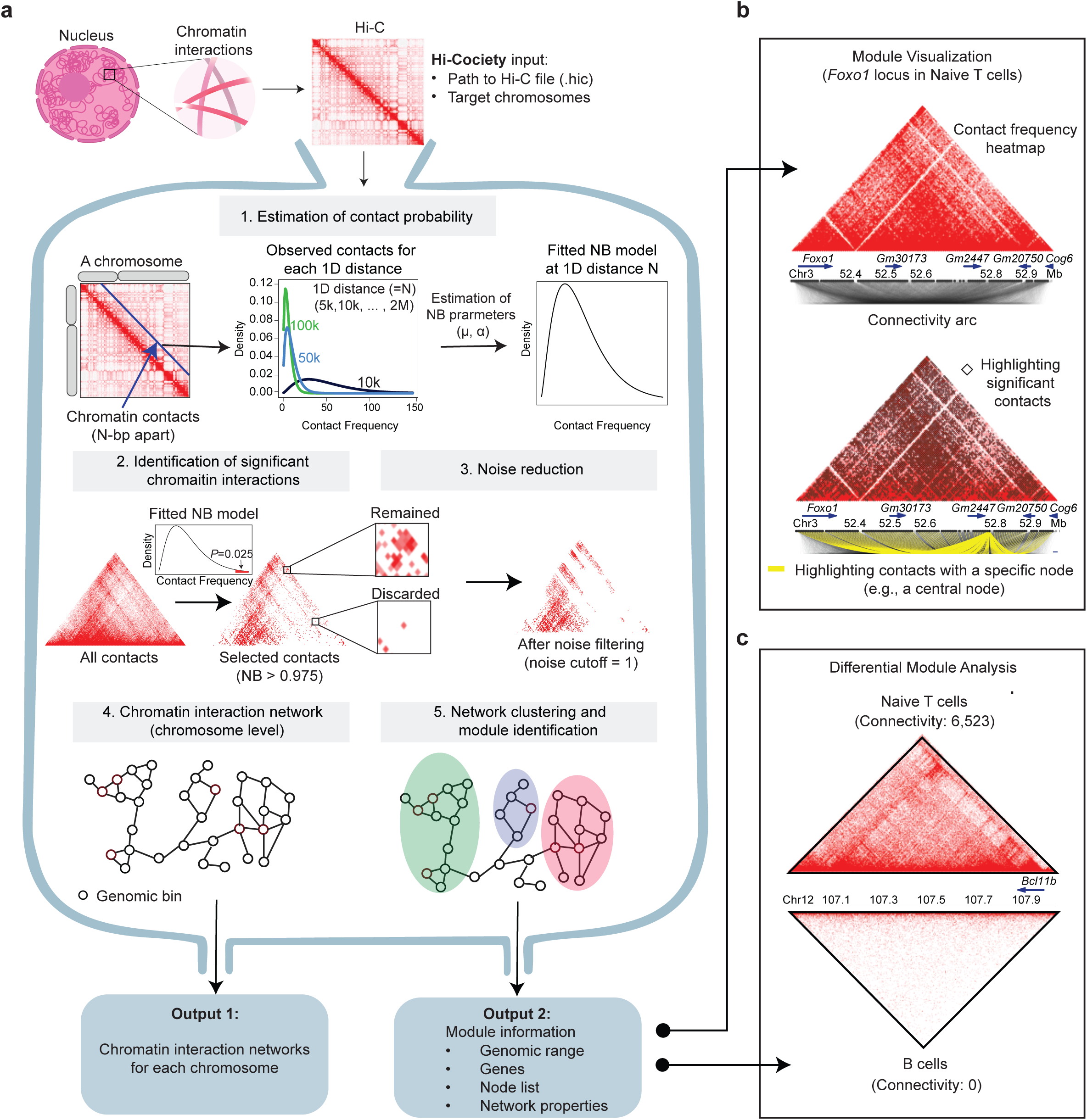
Network-based identification of significant chromatin interactions from Hi-C data. (a) Workflow of Hi-Cociety. The *hic2community* function in the HiCociety R package identifies chromatin interaction modules by first modeling contact frequencies from Hi-C data using negative binomial distribution to select significant interactions, and then filtering out noise. The resulting chromatin interaction network is then clustered into modules. The *ConnectivityDiff* function compares module connectivity between two cell types. (b) Visualization of a chromatin interaction module identified by Hi-Cociety, exemplified by a highly connected module centered around *Foxo1* in naïve CD4⁺ T cells. (Top) Hi-Cociety displays a contact frequency heatmap alongside an arc plot illustrating chromatin interactions. (Bottom) Significant chromatin contacts can be highlighted as black diamonds, and arc originating from a user-specified node (e.g., a central node) can be visualized. In this example, yellow arcs represent interactions stemming from a central node within the *Foxo1*-associated module. (c) Hi-Cociety provides a *ConnectivityDiff* function to identify differentially connected modules between two Hi-Cociety module objects. As an example, a comparison between naïve T cells and B cells reveals that the *Bcl11b* module exhibits the most distinct connectivity difference between the two cell types.

The genomic pairs with significant interactions are used to construct an interaction network using iGraph^7^. Subsequently, a label propagation algorithm^8^ is applied to group chromatin interactions as distinct modules. Key network properties—including connectivity, transitivity, and eigenvector centrality—are then computed for each module. In parallel, Hi-Cociety generates a Hi-C contact heatmap annotated with significant interactions and visualized with interaction arcs for each module (Figure 1b). Furthermore, Hi-Cociety provides a function that accepts module objects from two Hi-C datasets and generates a comparison table based on differences in connectivity (Figure 1c). Together, Hi-Cociety is a statistical method implemented in R (https://cran.r-project.org/web/packages/HiCociety/index.html) which infers modules of interactions from Hi-C datasets.

### Hi-C network node degrees follow an exponential distribution

We first investigated whether networks inferred by Hi-C across different cell types share any common features. To this end, we applied Hi-Cociety to seven Hi-C datasets spanning multiple immune cell types, including B lymphocytes and T lymphocytes such as double-positive thymocytes^9^, naïve CD4⁺ T cells, CD4⁺ T helper 1 (Th1)^10^ and Th2 cells, as well as embryonic stem cells^11^ and cells from the olfactory nervous system^12^. We investigated the distribution of node degree for each network, which is the number of interactions across genomic regions. We found that the node degree for each cell type followed an exponential distribution (Figure 2a). Most nodes showed relatively low connectivity, while a small portion of nodes were hyperconnected and displayed significantly high degree of connectivity (Figure 2a). Multi-enhancer hubs are defined as genomic domains carrying multiple enhancers interacting in three-dimensional space and regulating gene expression^1,10,13^. We next assessed if the highly connected nodes predicted by Hi-Cociety contain genomic regions scored as super-enhancers. Super-enhancers are characterized by their high density of active enhancers and their ability to drive the expression of genes critical for cell identity^14^. We compared the node degrees of two groups of nodes: those with super-enhancers and those without super-enhancers. Our analysis revealed that nodes containing super-enhancers exhibit significantly higher interactions with other nodes (Figure 2b). This finding confirms highly connected modules based on Hi-Cociety predictions contain active enhancers.

**Figure 2.**
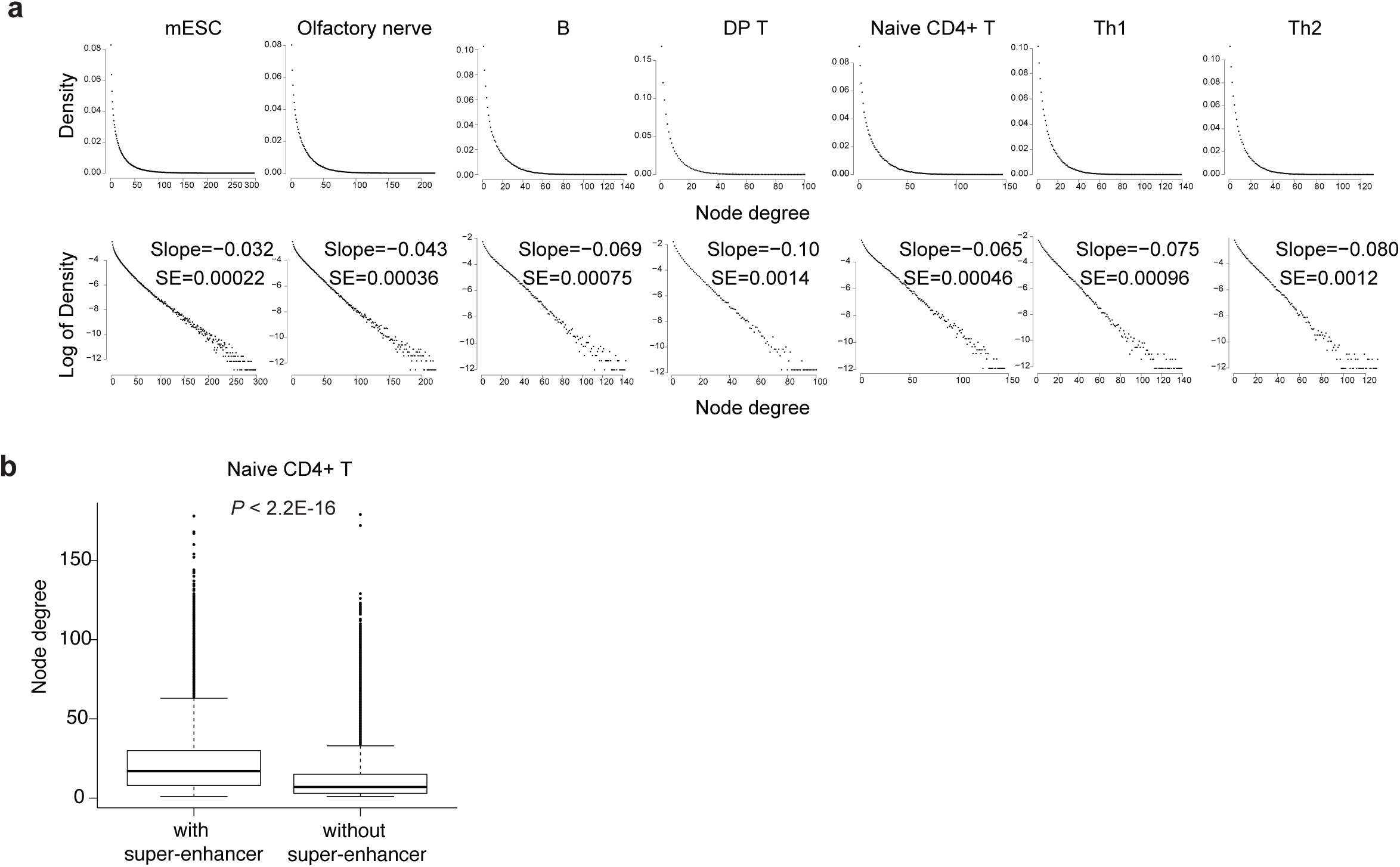
Network properties of Hi-Cociety-inferred significant chromatin interaction nodes. (a) Node degree distributions of significant chromatin interactions across seven independent datasets. (b) Comparison of node degree distributions in naïve CD4^+^ T cells between nodes harboring super-enhancers and those that do not.

### Highly connected chromatin interaction modules tend to be epigenetically active

We next examined the relationship between module connectivity and gene regulation in CD4⁺ T cells. Genes within the most highly connected modules were predominantly transcription factors with established roles in T cell biology (Figure 3a). These highly connected modules also ranked among the top in capturing subtype-specific signature genes, including *Bcl11b* in naïve CD4⁺ T cells and *Id2* in Th1/Th2 cells (Figure 3b, Table S1). Moreover, genes within the five most connected modules for each T cell subtype exhibited high expression levels (Figure S2a). To systematically explore the relationship between chromatin connectivity (Hi-C) and transcriptional activity, we analyzed bulk RNA-seq data from each T cell subtype (Figure 3c). We compared the top 100 highly connected chromatin modules ("top modules") with 100 randomly selected modules and assessed the expression levels of the most highly expressed genes within modules. Across all T cell subsets, genes within the top modules exhibited significantly higher expression than those in random modules (Figure 3c; naïve CD4⁺ T cells: *P* = 2.7 × 10⁻⁶; Th1: *P* = 1.2 × 10⁻⁴; Th2: *P* = 3.6 × 10⁻⁸). Consistent with this, chromatin accessibility and enhancer activity analyses revealed that highly connected modules were also more accessible and displayed elevated histone acetylation (Figure S2b–c). Together, these findings demonstrate that Hi-Cociety reliably detects high chromatin connectivity which is associated with increased epigenetic activity—marked by greater histone acetylation, enhanced accessibility, and higher gene expression.

**Figure 3.**
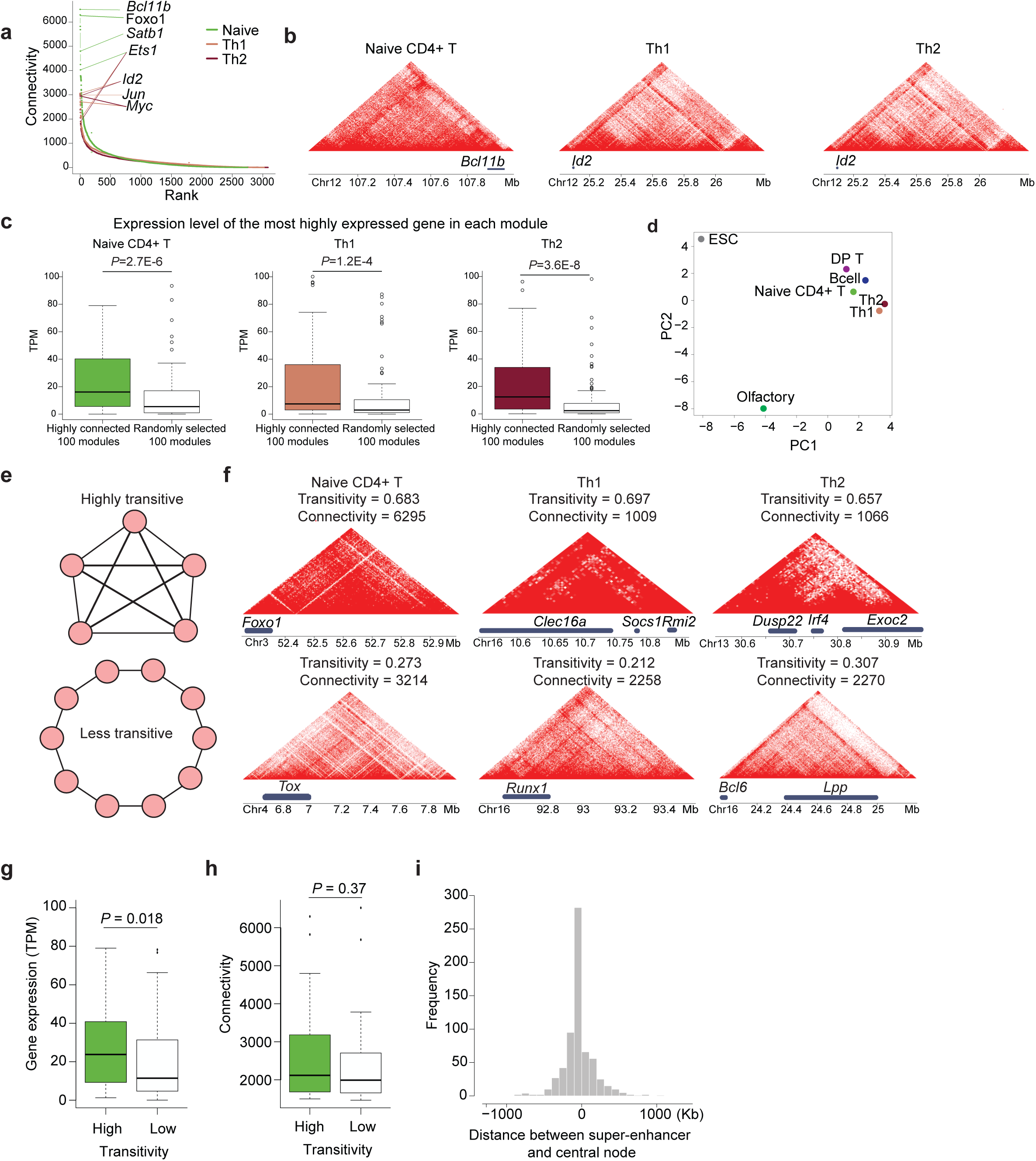
Network connectivity, transitivity, and centrality are closely associated with transcriptional outputs. (a) Distribution of network connectivity across chromatin interaction modules in naïve CD4⁺ T cells (green), Th1 cells (beige), and Th2 cells (brown), highlighting key genes. (b) Representative examples of highly connected modules in each T cell subtype. (c) Comparison of the expression levels of the most highly expressed genes between highly connected modules and randomly selected modules. (d) Principal component analysis (PCA) was performed on a binary matrix indicating whether each of the top 100 highly connected modules from each of the seven cell types overlapped with the merged set of all modules. (e) Schematic representations of modules with high (top) and low (bottom) transitivity; nodes represent genomic bins. (f) Example modules with high (top) and low (bottom) transitivity. (g–h) Comparisons of (g) gene expression (TPM scale) and (h) connectivity between high-transitivity (green) and low-transitivity (white) modules among the top 100 most connected modules in naïve CD4⁺ T cells. (i) Distribution of linear genomic distances between central nodes of Th1 modules and their nearest super-enhancers.

We also found that highly connected modules are cell-type-specific. We collected the top 100 highly connected modules across all cell types. Principal component analysis based on the binary presence or absence of the loci from each cell type revealed substantial differences between lymphocytes (e.g., B cells and T cells) and non-lymphocytes (e.g., embryonic stem cells and olfactory nerve cells) (Figure 3d). Among the top 100 olfactory nerve system Hi-C modules, we discovered 18 modules that include 53 G protein-coupled receptor genes essential for odor detection^15^ (Hypergeometric *P*-value = 0.030, Table S2). The highly connected loci were relatively similar among the closely related T cell subsets-naïve, Th1 and Th2 CD4^+^ T cells (Figure 3d). Thus, Hi-Cociety predicted highly connected modules corresponding to transcriptionally active genomic domains.

### Highly transitive modules exhibit increased transcriptional activity

In graph theory, a concept originated from social networks is called “transitivity”. This network feature is defined as the likelihood that relationships in a network are transitive, meaning that if node A is connected to node B, and node B is connected to node C, then there is a heightened probability that node A is also directly connected to node C. This property is often encapsulated in the concept of triadic closure, where a triad of nodes tends to form a closed triangle of connections. Hence, transitivity is defined as the proportion of interconnected triplets relative to any possible triplet in a network^16^ (Figure 3e). We used this definition and calculated transitivity of all modules across our datasets. Many genes fell within highly transitive modules where some have been implicated in T cell biology (Figure 3f, Table S3). An example includes *Foxo1*, which demonstrates a case of highly transitive module in T cell subsets (Figure 3f). To evaluate the relationship between transitivity and gene expression, we selected the top 100 modules from naive CD4^+^ T cell Hi-C based on connectivity, divided them into high and low transitivity groups, and compared the expression levels of the most highly expressed gene within each module. This analysis showed that genes within more transitive modules are significantly more expressed than genes within less transitive modules (Wilcoxon rank sum *P*-value = 0.018; Figure 3g). Interestingly, the connectivity of two groups was not significantly different (Wilcoxon rank sum *P*-value = 0.37; Figure 3h). Together, critical T cell genes are located at genomic regions with different degrees of connectivity and transitivity.

### Central nodes are enriched with super-enhancers, transcription factors and disease-associated SNPs

In graph theory, a central node serves as a primary influencer or hub within a network. One measure for quantifying this influence is “eigenvector centrality”, which evaluates the importance of a node not only based on the number of its connections but also on the quality of those connections. Specifically, eigenvector centrality assigns relative scores to nodes by considering the principle that connections to highly influential (highly scored) nodes contribute more to a node’s centrality than connections to less influential nodes. This recursive measure ensures that nodes embedded within well-connected neighborhoods are recognized as more central, making it particularly useful for identifying key players in complex networks.

To explore genomic elements that exert the most significant influence within each module of connected loci, we identified eigenvector centrality nodes in Th1 cells (Table S4). Remarkably, highly connected modules in Th1 cells with high centrality scores contain genes with critical roles in T cell biology. The node with highest centrality is *Id2* followed by *Foxo1, Myc, Bcl6, Bcl11b, Ets1,* and *Runx1* (Table S4). We then calculated the 1D genomic distance between each module’s central node and the nearest super-enhancers. Among 652 Hi-C modules containing super-enhancers, we detected an overlap between the super-enhancer and the node with the highest eigenvector centrality in 195 modules (30%). This is further supported by the distribution of distances between super-enhancers and central nodes, which has a mean of zero (Figure 3i). Hence, centrality analysis based on highly connected modules using Hi-Cociety suggests key regulatory hubs in T cells.

We next assessed whether highly connected modules are enriched for transcription factor (TF) genes by analyzing node centrality. For each of the top 100 modules, we classified nodes with the highest centrality scores as high-centrality nodes and those with the lowest scores as low-centrality nodes. Strikingly, TF genes were significantly overrepresented among high-centrality nodes (22%) compared to low-centrality nodes (4%) (Figure S2d), suggesting that genomic loci encoding transcription factors tend to occupy central positions within these chromatin interaction networks.

Cell type–specific super-enhancers are frequently enriched for disease-associated single-nucleotide polymorphisms (SNPs) in many developmental contexts^14,17,18^. We next asked whether genetic variants linked to complex immune disorders are preferentially enriched within nodes of high centrality. To address this, we compared the number of SNPs located within nodes of high versus low eigenvector centrality, as previously defined. Using the GWAS Catalog^19,20^, we identified SNPs overlapping genomic regions in each group. High centrality nodes overlapped with 374 SNPs, primarily associated with autoimmune diseases and asthma, whereas low centrality nodes overlapped with only 48 SNPs, showing limited enrichment for immune-related traits (Figure S2e). Together, these results indicate that highly central nodes are enriched for super-enhancers, transcription factor gene promoters, and disease-associated variants, suggesting that their central roles in immune gene regulation.

### Hi-Cociety can detect differential connectivity between experimental conditions

A core analytical capability of Hi-Cociety is its ability to detect genomic regions with significant differences in chromatin connectivity across experimental conditions. To evaluate this feature, we conducted three comparative analyses representing varying degrees of ontological similarity between cell types: B lymphocytes versus T lymphocytes (distinct lineages), naïve CD4^+^ T cells versus CD4^+^ Th1 cells (related subtypes), and wildtype Th1 cells versus super-enhancer-deleted Th1 cells (genetic perturbation). To define differential modules, we first identified highly connected chromatin regions in each cell type by selecting a connectivity threshold where the slope of the normalized connectivity curve was closest to 1. We then assessed the conservation of these modules in the comparison cell type and calculated fold changes in connectivity. Modules exhibiting a fold change greater than 1.3 were classified as cell-type–specific. As an initial test case, we compared naïve follicular B cells with naïve CD4^+^ T cells. Hi-Cociety identified 325 differential modules—71 specific to B cells and 254 specific to T cells (Tables S5–S6). Notably, the most B cell–specific module contained *Mef2c*, which showed 652 interactions in B cells and no detectable connectivity in T cells. *MEF2C* is known to cooperate with EBF-1 to activate B cell–specific transcriptional programs, directing lymphoid progenitors toward the B cell lineage^21^. Conversely, the most T cell–specific module included *Bcl11b*, which displayed 6,523 interactions in T cells and none in B cells. BCL11B plays an essential role in T cell development by suppressing innate immune genes and sustaining T cell receptor signaling components^22^. In both cases, increased chromatin connectivity strongly correlated with elevated gene expression in the respective cell types (Figure 4a), demonstrating Hi-Cociety’s ability to uncover functionally relevant structural differences.

**Figure 4.**
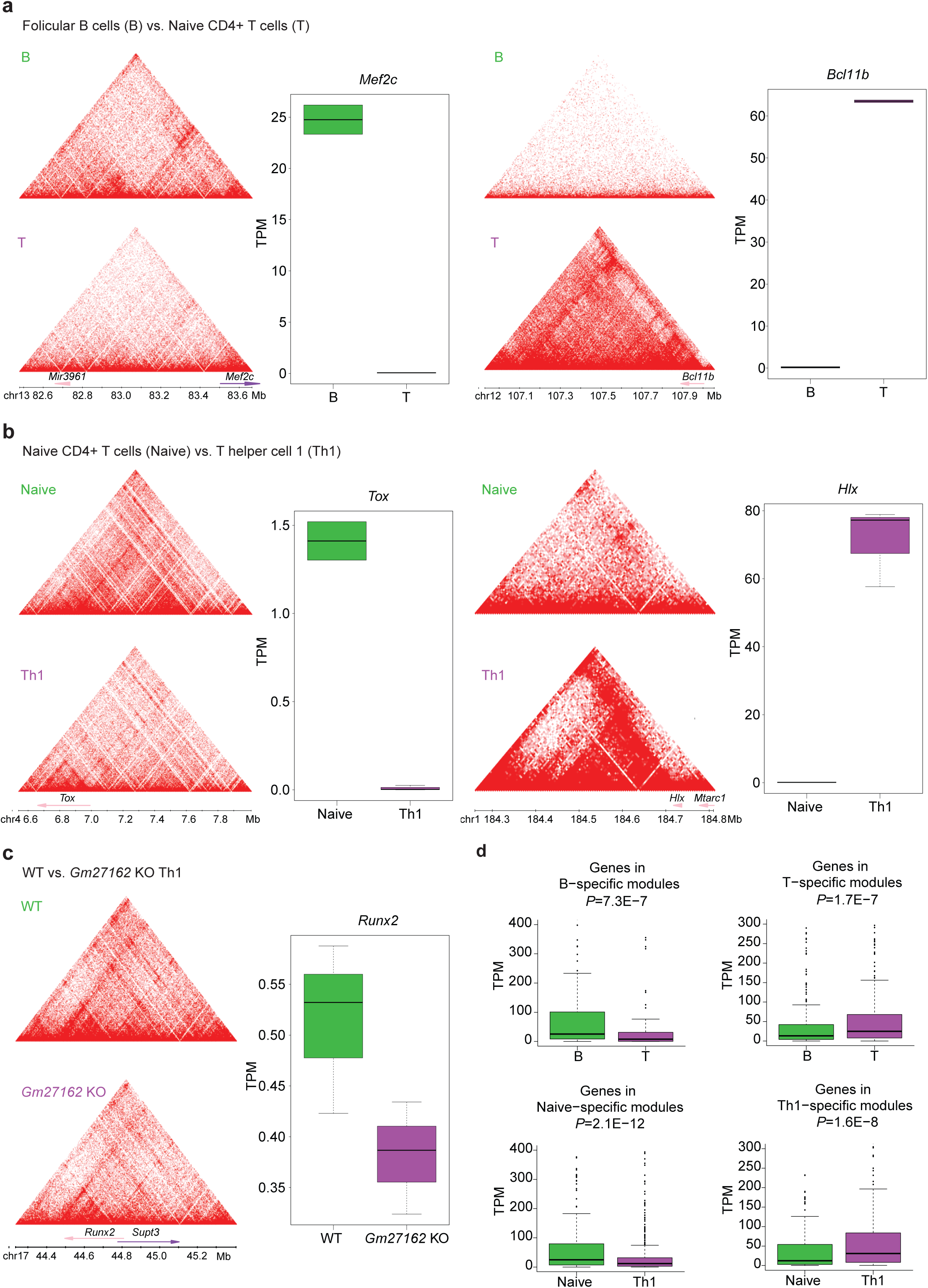
Hi-C modules with differential connectivity are associated with condition-specific gene expression. (a–c) Examples of modules showing differential connectivity and associated gene expression profiles: (a) Follicular B cells versus naïve CD4⁺ T cells, highlighting *Mef2c* (B cell-specific) and *Bcl11b* (T cell-specific). (b) Naïve CD4⁺ T cells versus Th1 cells, highlighting *Tox* (naïve T cell-specific) and *Hlx* (Th1-specific). (c) Wild-type versus *Gm27162*-deleted Th1 cells, highlighting *Runx2* (wild-type-specific). (d–e) Comparisons of maximum gene expression within condition-specific modules: (d) Follicular B cells versus naïve CD4⁺ T cells. (e) Naïve CD4⁺ T cells versus Th1 cells.

We next applied Hi-Cociety to compare chromatin modules between naïve CD4^+^ T cells and CD4^+^ Th1 cells. We observed 189 specific interactions in naïve T cells and 148 specific interactions in Th1 cells (Tables S7–S8). This finding suggests that even closely related T cell subsets possess distinct transcriptional and epigenetic architectures, corroborating previous reports indicating large-scale chromatin changes during T helper cell differentiation^23,24^. Among the most striking differences was the naïve CD4^+^ T cell-specific module containing *Tox*, a key regulator of CD4^+^ T cell lineage commitment^25^. This region exhibited dramatically higher connectivity in naïve cells (3,214 interactions) compared to Th1 cells (140 interactions), correlating with exclusive expression of *Tox* in naïve CD4^+^ T cells (Figure 4b). Conversely, one of the most pronounced Th1-specific modules included *Hlx* (ranked 2nd), which showed 18-fold stronger connectivity in Th1 cells (1,035 interactions) than in naïve cells (57 interactions). *Hlx* drives *Ifng* expression in differentiating helper T cells^26^ and forms a robust topologically associating domain (TAD) in Th1 cells, anchored at the *Hlx* locus. In naïve CD4^+^ T cells, this structural organization was markedly weaker, consistent with the significantly lower *Hlx* expression observed (Figure 4b). These results demonstrate how Hi-Cociety captures cell-type-specific chromatin architectures that underlie functional gene regulation, even between closely related immune cell subsets.

We also compared Hi-C of Th1 cells with another group of Th1 cells wherein the super-enhancer near *Ets1*, referred to as *Gm27162*, was genetically ablated. The small genetic deletion caused 69 wild type Th1-specific modules but no change in *Gm27162*-deleted Th1 cells (Table S9). The module with the most pronounced differential connectivity included *Runx2* (ranked 7th), a known target of ETS1^27^ with connectivity values of 1,075 in wild type and 493 *Gm27162*-deleted Th1 (Figure 4c). Although both wild type and *Gm27162*-deleted Th1 cells exhibited stripes at this locus, the intensity was much lower in the *Gm27162-*deleted Th1 cells. Concurrent with this finding, there was a notable decrease in *Runx2* gene expression observed in the *Gm27162*-deleted Th1 cells (Figure 4c).

To examine the link between connectivity and gene expression, we systematically analyzed the expression levels of genes located on B cell-specific and T cell-specific modules. Genes located on B cell-specific modules exhibited significantly higher expression in B cells (Wilcoxon rank sum *P*-value=7.3 × 10^⁻7^), whereas genes located on T cell-specific modules displayed significantly higher expression in T cells (Wilcoxon rank sum *P*-value=1.7 × 10^⁻7^) (Figure 4d). A similar pattern was observed for naïve CD4^+^ T cell-specific genes and Th1-specific genes (Figure 4d). Collectively, Hi-Cociety can effectively identify differential chromatin interaction modules for two cell types with varying levels of dissimilarity.

### Genetic deletion at the primary impact site *in silico* shows a greater impact when the module exhibits higher connectivity and lower transitivity

To examine the effect of genetic perturbation in genomic regions with distinct network properties such as connectivity and transitivity, we next performed *in silico* genetic perturbation using C.Origami^28^. C.Origami is a multimodal deep learning model that predicts cell-type-specific chromatin organization *de novo* from DNA sequence, CTCF binding, and chromatin accessibility data. Beyond modeling baseline chromatin structure, C.Origami can also simulate the effects of genetic deletions on local chromatin architecture. To validate its predictive power, we used experimental data following *Gm27162* deletion at the *Ets1-Fli1* locus in T cells^10^. We compared insulation scores predicted by C.Origami after *Gm27162* deletion to experimental Hi-C measurements in *Gm27162^-/-^* T cells (Figure S3a). The comparison revealed a strong correlation in the insulation scores between predicted and observed chromatin structures (R = 0.864, P < 2.2 × 10⁻¹⁶). Next, we systematically scanned chromosome 2 by performing *in silico* deletion of every 10-kb DNA segment and quantifying the resulting impact on chromatin organization within the surrounding ∼2-Mb region (Figure 5a). We applied Hi-Cociety to naïve CD4⁺ T cells and identified 205 chromatin modules on chromosome 2, 32 of which exhibited high connectivity (Figure 5b). We next stratified the modules by their connectivity and assessed the maximum impact scores within groups, which reflect the level of changes in C.Origami predictions of 3D chromatin folding following systematic deletion of 10-kb sequences. We found that highly connected modules were more susceptible to genetic perturbation (Figure 5c, S3b). We also stratified modules according to their transitivity but found that highly transitive modules were less susceptible to genetic perturbation (Figure 5d, S3c). Together, these findings suggest that distinct topological features of chromatin modules, such as connectivity and transitivity, differentially influence their sensitivity to genetic perturbation.

**Figure 5.**
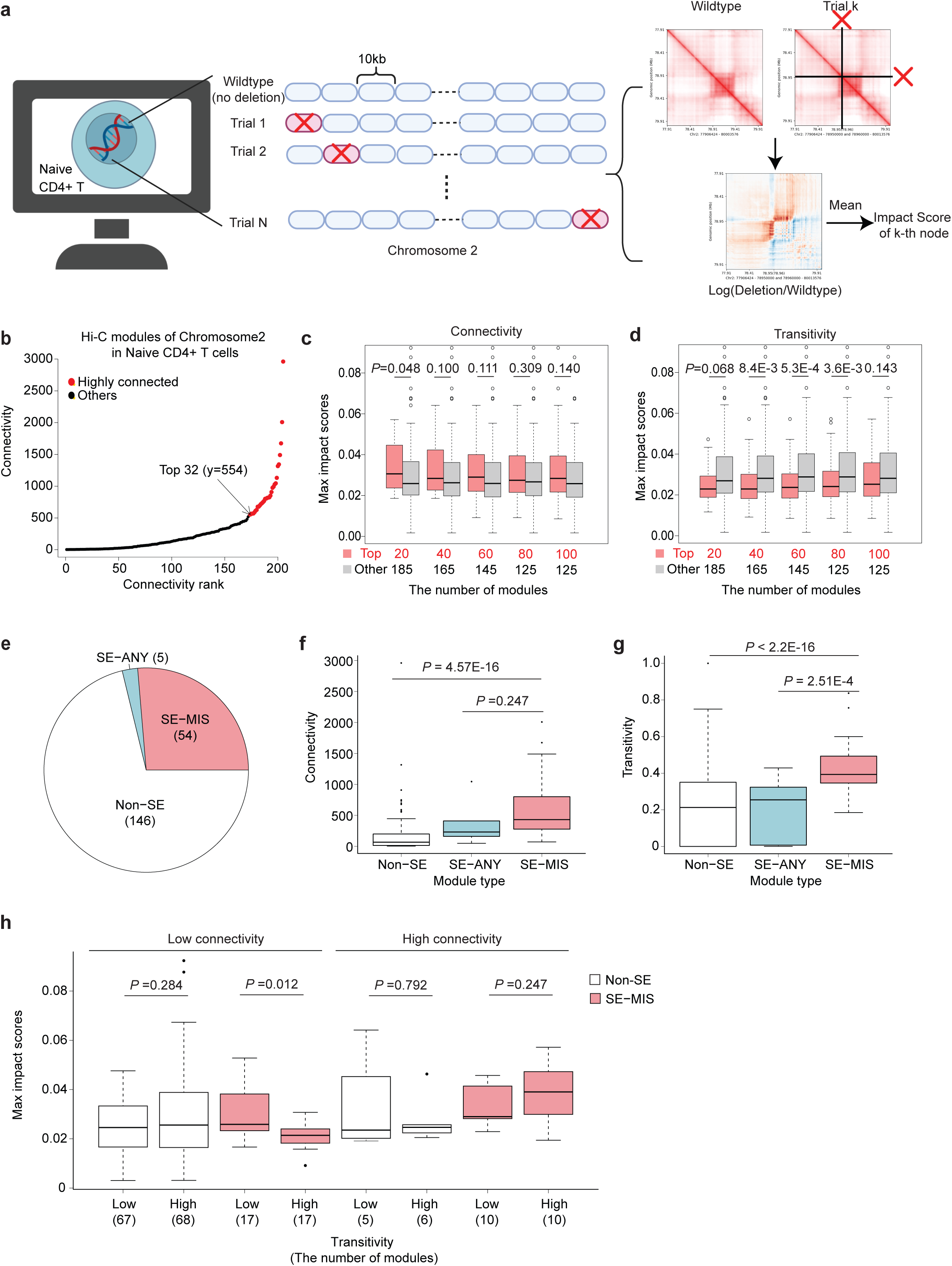
*In silico* perturbation analysis reveals that 3D chromatin reorganization depends on connectivity, transitivity, and super-enhancer presence. (a) *In silico* perturbation of mouse chromosome 2 (mm10) using C.Origami. Each 10-kb genomic bin was systematically deleted, and the impact score for each bin was calculated as the average log-fold change in contact frequencies relative to the wild-type prediction. (b) Connectivity distribution of modules on chromosome 2 in naïve CD4⁺ T cells, with highly connected modules highlighted in red. (c–d) Comparisons of maximum impact scores stratified by (c) connectivity and (d) transitivity. (e) Number of modules categorized based on super-enhancer presence. (f–g) Comparisons of (f) connectivity and (g) transitivity across modules stratified by super-enhancer type. (h) Comparison of maximum impact scores across modules grouped by connectivity, super-enhancer status, and transitivity.

### *In silico* deletion of primary impact sites disrupts local chromatin conformation

We next performed *in silico* genetic deletion analysis on modules containing super-enhancers. Among the 205 modules, 59 modules had at least one super-enhancer hence we refer to these modules as multi-enhancer hubs^10,13^. In 91.5% of these multi-enhancer hubs (54), a super-enhancer was overlapping either directly with the 10-kbp region showing the maximum impact score (primary impact site) or its immediate 10kbp neighborhood, defined as its first neighbor in the Hi-C network, suggesting genetic susceptibility of super-enhancers (Figure 5e). We stratified the modules into three categories: (1) SE-MIS, in which a super-enhancer either overlaps with the primary impact site or its first neighbor in the Hi-C network; (2) SE-ANY, where a super-enhancer is present but not meet the SE-MIS criteria; and (3) non-SE, modules lacking any super-enhancer. Connectivity and transitivity were both positively correlated with the presence of super-enhancers across modules (Figure 5f,g). To evaluate the combined effects of connectivity, transitivity, and the presence of super-enhancers at SE-MIS on maximum impact scores, we classified modules into eight groups based on these three factors (high/low connectivity, high/low transitivity, and super-enhancer presence). This analysis revealed that connectivity and transitivity jointly influenced maximum impact scores, particularly in the presence of super-enhancers (Figure 5h). Highly connected modules containing super-enhancers exhibited consistently higher maximum impact scores, regardless of their transitivity. This suggests that in highly connected modules, primary impact sites marked by high H3K27ac levels may act as spatial hubs that coordinate extensive interactions, and their removal leads to widespread topological reorganization. In contrast, for modules with low connectivity, we observed that transitivity became a significant determinant of the maximum impact score, but only in modules containing super-enhancers. Specifically, modules with both low connectivity and low transitivity exhibited significantly higher impact scores (Figure 5h), which may reflect the reduced presence of alternative acetylated loci in their first neighborhood in the Hi-C network (Figure S3e). This implies that when both connectivity and transitivity are low, primary impact sites may act as critical points of control over local chromatin topology, and their perturbation has a more pronounced effect. Conversely, when transitivity is high in low-connectivity modules with the presence of super-enhancer, the spatial proximity of multiple acetylated regions may buffer the loss of primary impact site, leading to lower maximum impact scores (Figure 5h).

Taken together, Hi-Cociety models 3D chromatin architecture as a network of interacting elements, revealing that network properties such as connectivity, transitivity, and centrality are closely associated with gene expression regulation. In-silico deletion analyses further showed that chromatin network modules containing super-enhancers are particularly vulnerable to genetic perturbations, with sensitivity increasing when connectivity is high or when connectivity and transitivity are low.

## Discussion

In this study, we introduced Hi-Cociety, a graph-based computational framework that leverages Hi-C data to infer 3D enhancer communities and chromatin interaction modules without relying on histone modification or chromatin accessibility data. By constructing a network of significant chromatin interactions and applying clustering algorithms, Hi-Cociety successfully identified spatially connected genomic regions that function as regulatory hubs. Our analysis revealed that highly connected chromatin modules are enriched for super-enhancers, transcription factors, and disease-associated SNPs, and are associated with increased transcriptional activity, chromatin accessibility, and histone acetylation. These findings highlight the importance of 3D chromatin organization in gene regulation and suggest that highly connected modules serve as key regulatory hubs that integrate multiple signaling inputs to ensure robust and precise control of lineage-specific transcriptional programs. Furthermore, Hi-Cociety’s ability to detect differential chromatin interaction patterns between cell types underscores its utility in identifying cell-type-specific regulatory architectures, providing new insights into the role of 3D chromatin organization in cellular differentiation and disease.

The *in silico* genetic perturbation analysis further demonstrated that highly connected modules with lower transitivity are more susceptible to local chromatin reorganization, particularly when they harbor super-enhancers. This suggests that the network properties of chromatin interaction modules, such as connectivity and transitivity, play a critical role in determining their resilience to genetic perturbations. Modules with high connectivity and low transitivity may act as critical regulatory hubs that are more sensitive to disruptions, potentially leading to significant changes in gene expression and cellular function. These findings have important implications for understanding the mechanisms underlying developmental disorders and diseases such as cancer, where disruptions in 3D chromatin organization can lead to aberrant gene expression. Overall, Hi-Cociety provides a powerful and scalable tool for studying 3D chromatin architecture, offering new avenues for exploring the role of enhancer communities in gene regulation and disease.

## Methods

### Hi-Cociety pipeline

Hi-Cociety is an R package that aims to find chromatin interaction modules and detect differential chromatin folding patterns from two sets of chromatin conformation capture measurements such as Hi-C and micro-C. To accomplish this, Hi-Cociety first captures various chromatin folding patterns within the genome conformation capture measures. Afterwards, it estimates the difference in connectivity of each module in two different datasets. The pipeline by which Hi-Cociety detects difference in chromatin folding patterns in two datasets can be described as follows:

(1) Loading of .hic file: The input files are formatted in .hic. Those files are loaded in R as a three-column data frame through the straw function in the strawr R package ^29^.
(2) Estimation of contact probability: The contact frequencies are then stratified based on the one-dimensional (1D) genomic distance. By default, 1D genomic distance is considered up to 2Mb. Next, size and mean (mu) parameters for each group of contact frequencies are estimated assuming the counts follow negative binomial distribution. Then the probability value of each count was calculated based on the estimated negative binomial parameters. By default, the counts of which probability values are equal or larger than 0.975 is considered to be significant.
(3) Deletion of noisy chromatin interaction: Given that significant contacts in regions with low contact density may potentially be noise, we opted to eliminate such contacts. To address this, we introduced a 5-pixel padding to significant contacts and decided to exclude any contact where the average contact frequency in the extended region is less than *N*. Here, *N* is user-defined, with a default value of 1.
(4) Conversion of Hi-C to network: For the remaining significant chromatin interactions, network object is created by utilizing *graph_from_data_frame* function in igraph R package^30^.
(5) Module detection: To find the chromatin interaction modules, the network is clustered using label propagation method or Louvain clustering implemented as *cluster_label_prop* or *cluster_louvain* function in iGraph R package^30^, respectively. The difference between label propagation method and Louvain clustering is the implementation of randomness. We used label propagation method for the data in this manuscript. All parameters are set as defaults. Modules of which number of nodes is more than 3 are retained.
(6) Comparison of module: This process is done with *ConnectivityDiff* function in HiCociety R package. This function takes as input Hi-C module objects for two distinct cell types, as derived from the ‘*hic2community*’ function. Initially, it computes connectivity of each module for counterpart cell type. Subsequently, the fold-change in connectivity between two cell types is calculated. Cell-type-specific modules are identified using a connectivity cutoff determined by the ‘*getElbowPoint’* function in the HiCociety R package. This function normalizes the x-axis (rank of modules by connectivity), and the y-axis (connectivity) to a range of [0,1], and then identifies the point where the slope is closest to 1.

### Collection of public Hi-C data

Hi-C data of mouse embryonic stem cells (mESC)^11^, olfactory nerve system^31^ and B cell lymphoma^32^ were obtained from 4DNucleome data portal (4DNFI4OUMWZ8.hic, 4DNFIXKC48TK.hic, and 4DNFI8KBXYNL.hic, respectively). Hi-C data of DP T cells^9^, wildtype and Gm27162-deleted Th1 cells^10^ were obtained from Gene Expression Omnibus^33,34^ (GSE178345 and GSE211176, respectively).

### Collection of public RNA-seq data

RNA-seq of mouse naïve CD4^+^ T cells, Th1 cells and *Gm27162*-deleted Th1 cells were obtained from the Gene Expression Omnibus^33,34^ (GSE211177).

### Node degree distribution

The node degree of networks from seven Hi-C datasets were calculated using *degree.distribution* function from *igraph* R package^7^. Similarly, log of node degree density plot was estimated by taking log (natural constant as base) to the density values.

### Super-enhancer detection

Super-enhancers in Th1 and naïve CD4^+^ T cells were identified using H3K27ac ChIP-seq as previously described^14^. In summary, H3K27ac peaks were merged across each 12,500 bp genomic bin, and ChIP-seq occupancy was ranked in ascending order. The occupancy values were then rescaled to a range between 0 and 1. The ranked ChIP-seq occupancy showed a rapid increase at a point where the slope of the curve reached 1. Super-enhancers were defined as regions where ChIP-seq occupancy exceeded this point.

### Connectivity, transitivity and centrality estimation

The connectivity, transitivity and centrality of a network or module were estimated using function *length(E())*, *transitivity* and *authority.score* in iGraph R package^7^, respectively.

### Triangle heatmap alongside an arc plot

A triangle heatmap combined with an arc plot for a Hi-Cociety module is generated by utilizing *vizualizeModule* function in HiCociety R package, with connections to central nodes highlighted. For some triangle heatmaps, arc plots were removed, and gene tracks were enhanced using Adobe illustrator.

### Principal component analysis

For seven Hi-C datasets encompassing mouse embryonic stem cells, the mouse olfactory nerve system, B cells, DP T cells, naïve CD4^+^ T cells, Th1 cells, and Th2 cells, Hi-Cociety was executed using default parameters. Subsequently, the top 100 modules with the highest connectivity were chosen from each dataset, and corresponding bed data frames for genomic coordinates were constructed. These frames were then amalgamated to generate a binary table illustrating the intersection between the merged coordinates and those of each individual dataset. Principal component analysis was conducted on this table utilizing the *prcomp* function in the R base package^35^.

### Centrality and non-centrality nodes

Here, we defined the centrality and non-centrality nodes of a module as those with the highest and lowest scores from the eigenvector centrality assessment. The centrality and non-centrality nodes were compared in terms of the overlap of genomic range with the transcription factor gene and SNP occupancy as following.

### Transcription factor gene coordinate information

The coordinates of mouse transcription factor genes were acquired from the Homo sapiens Comprehensive Model Collection (HOCOMOCO) version 11, which supplies binding models for transcription factors encompassing 680 human and 453 mouse transcription factors^36^. We compared the occupancies of transcription factor genes at centrality and the non-central nodes, and the frequency is illustrated as pie chart in Figure S2d.

### Single nucleotide polymorphism (SNP) analysis

To compare the number and types of SNPs within centrality and non-centrality nodes, data from a recent genome-wide association study (GWAS) was downloaded from the NHGRI-EBI GWAS catalog (version 2023-02-15)^19,20^. We then mapped the human genomic coordinates corresponding to the mouse centrality and non-centrality nodes, respectively using LiftOver^37^. The overlap between centrality and non-centrality nodes with the SNP loci was evaluated using *findOverlaps* function in *GenomicRanges* R package^38^.

### *In silico* genetic perturbation analysis

We conducted an *in silico* genetic deletion analysis using C.Origami^28^ which predicts local chromatin folding change following the deletion of a genomic locus by utilizing a multimodal deep neural network. It consists of two steps: (1) model training and (2) prediction of the deletion effect on the local 3D chromatin conformation. For model training, we used *‘corigami-train’* function and provided the sequence of mouse mm10 (in FASTA format), Hi-C data of mouse DP T cells (in .npz format), and CTCF ChIP-seq and ATAC-seq data (in bigwig format) of the same cell type. The Hi-C data were converted from .cool format to .npz using the ’cool2npy.py’ script provided by C.Origami. The signals from CTCF ChIP-seq and ATAC-seq were used after taking the logarithm of their values. Next, to systematically delete every 10kb within chromosome 2 of naïve CD4^+^ T cells, we utilized ‘*corigami-screen’* function. Here, we provided the trained model, the path to the sequence file and the bigwig file of ATAC-seq and CTCF ChIP-seq data for the mouse naïve CD4^+^ T cells. Additional options such as screen-start=10,000,000, screen-end=179,913,224, perturb-width=10,000, step-size=10,000 and padding=0 were set. The resulting bedgraph file provides the impact score of each deletion which is defined as the average log of fold change of pixels (deletion/wildtype) within predicted Hi-C map covering ∼2Mb region. Among them we selected maximum impact score for each of 205 modules from chromosome 2 of mouse naïve CD4^+^ T cells predicted by Hi-Cociety. Then we investigated the effects of super-enhancer, connectivity and transitivity on the maximum impact score as shown in Figure 5.

## Data and code availability

HiCociety R package is available on CRAN and the data and code are available in the author’s Github page: https://github.com/ysora/HiCociety. Tables of chromatin interaction modules from three T cell subtypes, along with the results of differential module analysis results are available at the following web site: https://sorayoon.shinyapps.io/HiCociety/.

## Author Contributions

Sora Yoon contributed to the conceptualization, methodology, software development, validation, formal analysis, investigation, data curation, writing of the original draft, review and editing of the manuscript, and visualization.

Golnaz Vahedi contributed to resources, supervision, project administration, funding acquisition, and review and editing of the manuscript.

## Acknowledgments

We thank helpful discussion with members of the Vahedi laboratory in particular Dr. Aditi Chandra and Atishay Jay. We thank the Faryabi laboratory in particular Tobias Friedrich for useful comments. This study was supported by the Burroughs Welcome Fund, the Chan Zuckerberg Initiative Award, and NIH U01 DK112217, R01AI168240, U01 DK127768, U01 DA052715 (G.V).

---

**Figure S1.**
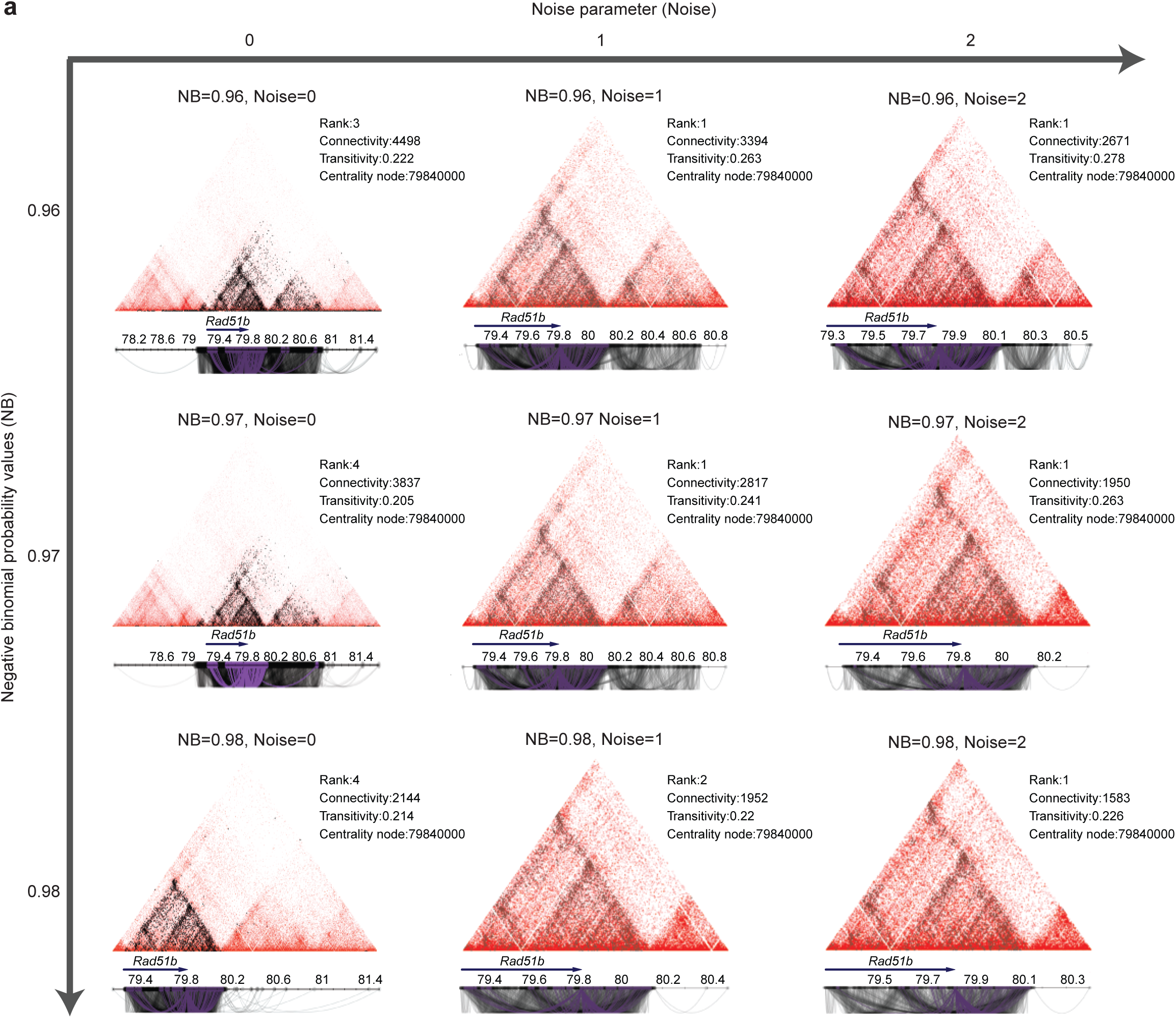
Parameter optimization for Hi-Cociety. (a) To suggest the optimal values for the negative binomial (NB) and noise parameters for defining chromatin modules, we tested various parameter combinations using Hi-C data with a fixed number of valid pairs. Specifically, we downsampled deeply sequenced large pre-B cell Hi-C data^6^ to 2 million valid pairs and evaluated module formation at the Rad51b locus under nine parameter combinations (NB = 0.96, 0.97, 0.98; noise = 0, 1, 2). As expected, higher NB values and stricter noise cutoffs led to the identification of denser modules. Notably, the difference between noise settings 0 and 1 was substantial, highlighting the critical importance of filtering out noisy chromatin interactions.

**Figure S2.**
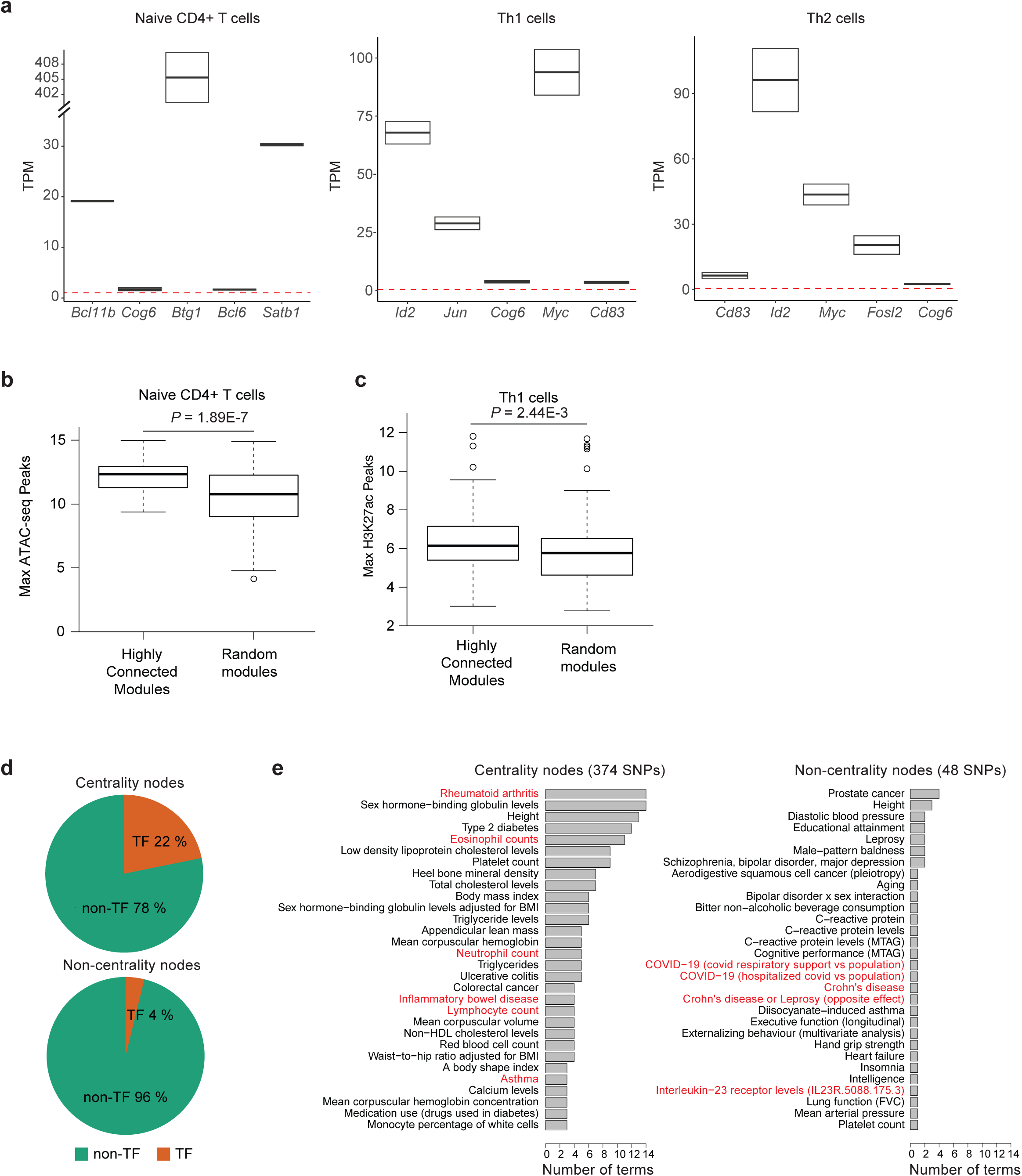
Connectivity of modules correlates with transcriptional and epigenetic activity. (a) Expression levels of representative genes from the five most highly connected modules in each of three T cell subtypes. The dashed red line indicates the median expression level of all genes with TPM > 0. (b–c) Comparisons of (b) maximum chromatin accessibility (ATAC-seq peaks) and (c) maximum histone acetylation (H3K27ac) levels between the 100 most highly connected modules and 100 randomly selected modules. (d) In Th1 cells, 22% of central nodes in the top 100 modules encode transcription factors, compared to only 4% of non-central nodes. (e) Central nodes harbor a greater number of SNPs associated with immune diseases compared to the least central nodes.

**Figure S3.**
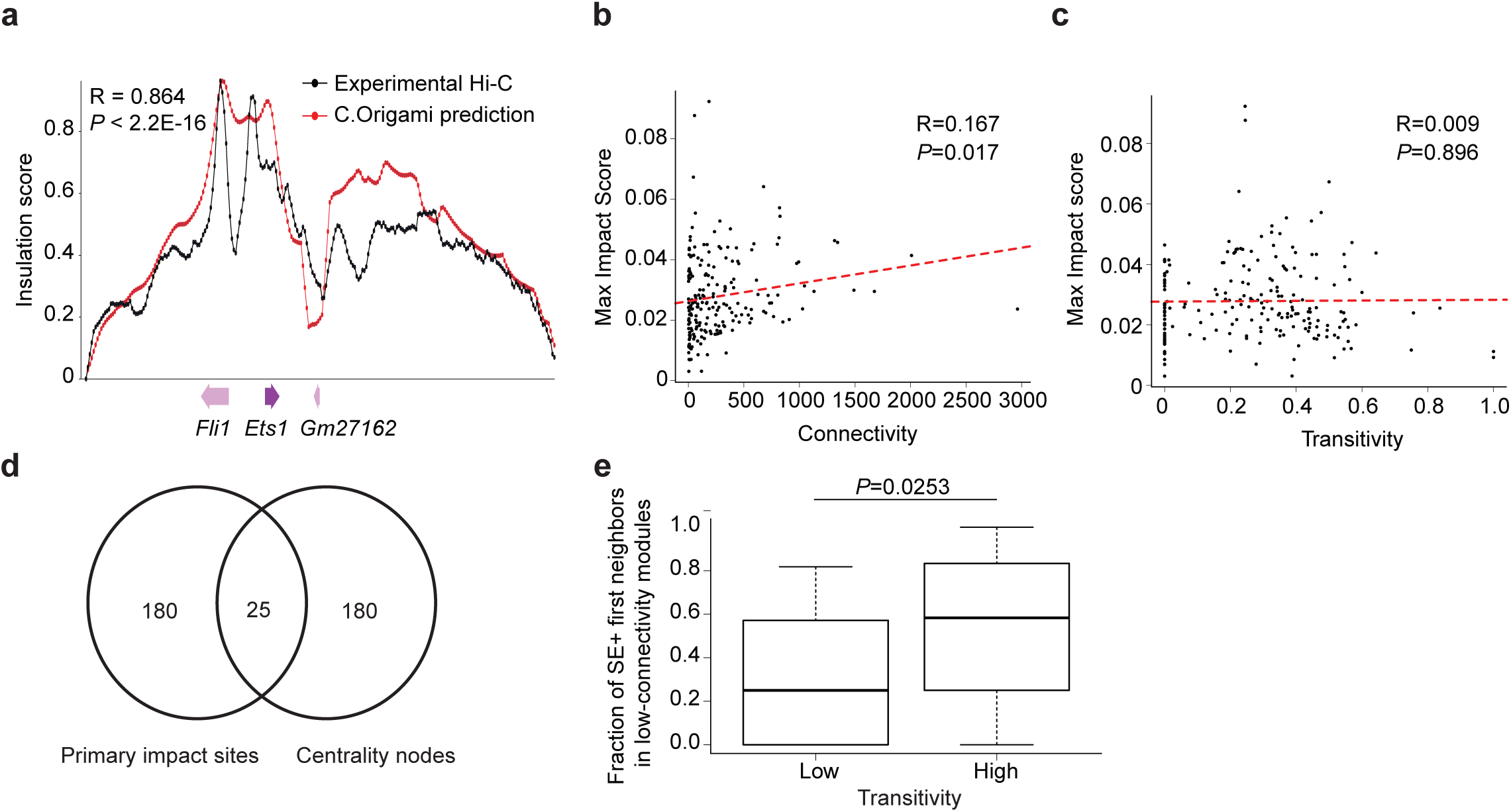
Validation of C.Origami predictions through in silico perturbation analysis. (a) Insulation scores from experimental Hi-C data of *Gm27162*-deleted naïve CD4⁺ T cells strongly correlate with C.Origami predictions at the *Fli1-Ets1* locus (R = 0.864). (b–c) Relationships between maximum impact score and (b) connectivity and (c) transitivity among modules on chromosome 2 in naïve CD4⁺ T cells. (d) Overlap between primary impact sites identified by C.Origami and central nodes in each module. (e) For lower-connectivity modules containing super-enhancer, the fraction of first-neighbor nodes that also harbor super-enhancers was compared between modules with high and low transitivity.

## References

1. Uyehara, C.M. & Apostolou, E. 3D enhancer-promoter interactions and multi-connected hubs: Organizational principles and functional roles. Cell Rep 42, 112068 (2023).

2. Di Giammartino, D.C. et al. KLF4 is involved in the organization and regulation of pluripotency-associated three-dimensional enhancer networks. Nat Cell Biol 21, 1179–1190 (2019).

3. Petrovic, J. et al. Oncogenic Notch Promotes Long-Range Regulatory Interactions within Hyperconnected 3D Cliques. Mol Cell 73, 1174–1190 e12 (2019).

4. Fasolino, M. et al. Genetic Variation in Type 1 Diabetes Reconfigures the 3D Chromatin Organization of T Cells and Alters Gene Expression. Immunity 52, 257–274 e11 (2020).

5. Zhao, J. & Faryabi, R.B. Spatial promoter-enhancer hubs in cancer: organization, regulation, and function. Trends Cancer 9, 1069–1084 (2023).

6. Hu, Y.G. et al. Lineage-specific 3D genome organization is assembled at multiple scales by IKAROS. Cell 186, 5269-+ (2023).

7. Csardi, G. & Nepusz;, T. The igraph software package for complex network research. InterJournal Complex Systems, 1695 (2006).

8. Raghavan, U.N., Albert, R. & Kumara, S. Near linear time algorithm to detect community structures in large-scale networks. Phys Rev E Stat Nonlin Soft Matter Phys 76, 036106 (2007).

9. Wang, W. et al. TCF-1 promotes chromatin interactions across topologically associating domains in T cell progenitors. Nat Immunol 23, 1052–1062 (2022).

10. Chandra, A. et al. Quantitative control of Ets1 dosage by a multi-enhancer hub promotes Th1 cell differentiation and protects from allergic inflammation. Immunity 56, 1451–1467 e12 (2023).

11. Bonev, B. et al. Multiscale 3D Genome Rewiring during Mouse Neural Development. Cell 171, 557–572 e24 (2017).

12. Monahan, K., Horta, A. & Lomvardas, S. LHX2- and LDB1-mediated trans interactions regulate olfactory receptor choice. Nature 565, 448–453 (2019).

13. Tsai, A., Alves, M.R. & Crocker, J. Multi-enhancer transcriptional hubs confer phenotypic robustness. Elife 8(2019).

14. Vahedi, G. et al. Super-enhancers delineate disease-associated regulatory nodes in T cells. Nature 520, 558–62 (2015).

15. Poll, B.G., Chen, L., Chou, C.L., Raghuram, V. & Knepper, M.A. Landscape of GPCR expression along the mouse nephron. Am J Physiol Renal Physiol 321, F50–F68 (2021).

16. Batjargal, B. Network triads: Transitivity, referral and venture capital decisions in China and Russia. Journal of International Business Studies 38, 998–1012 (2007).

17. Hnisz, D. et al. Super-enhancers in the control of cell identity and disease. Cell 155, 934–47 (2013).

18. Whyte, W.A. et al. Master transcription factors and mediator establish super-enhancers at key cell identity genes. Cell 153, 307–19 (2013).

19. Cerezo, M. et al. The NHGRI-EBI GWAS Catalog: standards for reusability, sustainability and diversity. Nucleic Acids Res 53, D998–D1005 (2025).

20. Sollis, E. et al. The NHGRI-EBI GWAS Catalog: knowledgebase and deposition resource. Nucleic Acids Research 51, D977–D985 (2023).

21. Kong, N.R., Davis, M., Chai, L., Winoto, A. & Tjian, R. MEF2C and EBF1 Co-regulate B Cell-Specific Transcription. PLoS Genet 12, e1005845 (2016).

22. Cismasiu, V.B. et al. BCL11B participates in the activation of IL2 gene expression in CD4+ T lymphocytes. Blood 108, 2695–702 (2006).

23. Vahedi, G. et al. STATs shape the active enhancer landscape of T cell populations. Cell 151, 981–93 (2012).

24. Ciofani, M. et al. A validated regulatory network for Th17 cell specification. Cell 151, 289–303 (2012).

25. Niu, H. & Wang, H. TOX regulates T lymphocytes differentiation and its function in tumor. Front Immunol 14, 990419 (2023).

26. Zheng, W.P. et al. Up-regulation of Hlx in immature Th cells induces IFN-gamma expression. J Immunol 172, 114–22 (2004).

27. Xia, W.Q., Han, X. & Wang, L. E26 transformation-specific 1 is implicated in the inhibition of osteogenic differentiation induced by chronic high glucose by directly regulating Runx2 expression. Journal of Biomedical Research 36, 39–47 (2022).

28. Tan, J. et al. Cell-type-specific prediction of 3D chromatin organization enables high-throughput in silico genetic screening. Nat Biotechnol 41, 1140–1150 (2023).

29. Durand, N. & Shamim, M. strawr: Fast Implementation of Reading/Dump for .hic Files. (2023).

30. Csárdi, G. et al. igraph: Network Analysis and Visualization in R_. (2024).

31. Bashkirova, E. & Lomvardas, S. Olfactory receptor genes make the case for inter-chromosomal interactions. Current Opinion in Genetics & Development 55, 106–113 (2019).

32. Vian, L. et al. The Energetics and Physiological Impact of Cohesin Extrusion. Cell 175, 292–294 (2018).

33. Edgar, R., Domrachev, M. & Lash, A.E. Gene Expression Omnibus: NCBI gene expression and hybridization array data repository. Nucleic Acids Res 30, 207–10 (2002).

34. Barrett, T. et al. NCBI GEO: archive for functional genomics data sets--update. Nucleic Acids Res 41, D991–5 (2013).

35. Team, R.C. R: A language and environment for statistical computing. (ed. Computing, R.F.f.S.) (Vienna, Austria, 2021).

36. Kulakovskiy, I.V. et al. HOCOMOCO: towards a complete collection of transcription factor binding models for human and mouse via large-scale ChIP-Seq analysis. Nucleic Acids Res 46, D252–D259 (2018).

37. Hinrichs, A.S. et al. The UCSC Genome Browser Database: update 2006. Nucleic Acids Res 34, D590–8 (2006).

38. Lawrence, M. et al. Software for computing and annotating genomic ranges. PLoS Comput Biol 9, e1003118 (2013).

